# *Toxoplasma gondii* Toxolysin 4 contributes to efficient parasite egress from host cells

**DOI:** 10.1101/2021.05.13.444112

**Authors:** My-Hang Huynh, Marijo S. Roiko, Angelica O. Gomes, Ellyn N. Schinke, Aric J. Schultz, Swati Agrawal, Christine A. Oellig, Travis R. Sexton, Jessica M. Beauchamp, Julie Laliberté, Komagal Kannan Sivaraman, Louis B. Hersh, Sheena McGowan, Vern B. Carruthers

## Abstract

Egress from host cells is an essential step in the lytic cycle of *T. gondii* and other apicomplexan parasites; however, only a few parasite secretory proteins are known to affect this process. The putative metalloproteinase Toxolysin 4 (TLN4) was previously shown to be an extensively processed microneme protein, but further characterization was impeded by the inability to genetically ablate *TLN4*. Herein we show that TLN4 has the structural properties of an M16 family metalloproteinase, that it possesses proteolytic activity on a model substrate, and that genetic disruption of TLN4 reduces the efficiency of egress from host cells. Complementation of the knockout strain with the TLN4 coding sequence significantly restored egress competency, affirming that the phenotype of the Δ*tln4* parasite was due to the absence of TLN4. This work identifies TLN4 as the first metalloproteinase and the second microneme protein to function in *T. gondii* egress. The study also lays a foundation for future mechanistic studies defining the precise role of TLN4 in parasite exit from host cells.

**IMPORTANCE:** After replicating within infected host cells, the single celled parasite *Toxoplasma gondii* must rupture out of such cells in a process termed egress. Although it is known that *T. gondii* egress is an active event that involves disruption of host-derived membranes surrounding the parasite, very few proteins that are released by the parasite are known to facilitate egress. In this study we identify a parasite secretory protease that is necessary for efficient and timely egress, thus laying the foundation for understanding precisely how this protease facilitates *T. gondii* exit from host cells.

## INTRODUCTION

Egress by apicomplexan parasites including *Toxoplasma gondii* and malaria parasites (*Plasmodium spp.*) is the critical last step in the lytic cycle. Egress liberates the parasite for infection of new cells and releases host cell cytosolic contents, which can activate an inflammatory immune response (1). Inflammation is a hallmark of toxoplasmosis, with associated ocular, neural, cardiac, or respiratory disease, which is especially severe in immunodeficient or congenitally infected individuals. Severe inflammation during toxoplasmosis is linked to the parasite genotype, with highly virulent strains being associated with worse outcome, especially for ocular and congenital infection (2).

Recent focus on egress is elucidating signaling pathways and associated secondary messengers, thus providing a broader understanding of the intricate interplay between the signals involved in this complex cascade of events (recently reviewed in (3). Such studies have solidified roles for cyclic GMP (cGMP) and calcium to respectively stimulate protein kinase G (PKG)(4, 5) and calcium-dependent protein kinases (CDPKs)(4, 6–8), among other targets. Calcium signaling results in the activation of parasite motility and the discharge of apical secretory granules termed micronemes (9–12). While the importance of microneme proteins such as transmembrane adhesins that connect with the actin-myosin motor system to drive gliding motility and active cell invasion have been established, microneme proteins also contribute to active egress via their role in the formation of pores such as those created by Perforin-Like Protein 1 (PLP1) (13–15). Gene knockout studies suggest that PLP1 pore formation disrupts the parasitophorous vacuole membrane (PVM), which encases parasites during intracellular replication. PLP1 deficient parasites are delayed or fail in egress and show a marked loss of virulence in infected mice, implying a link between efficient egress and virulence. Although several proteins released from parasite dense granules (calcium independent secretory organelles released during parasite replication) have also been implicated in egress (16–20), PLP1 is the only microneme protein known to directly function in egress to date.

Previous work identified a putative metalloprotease Toxolysin 4 (TLN4, TGME49_206510) in a proteomic screen of *Toxoplasma* secretory products released by extracellular tachyzoites (21). A subsequent study showed that TLN4 is a microneme protein that undergoes extensive proteolytic processing and potentially contributes to parasite fitness, based on loss of TLN4 deficient parasites from a mixed population of parasites transfected with a knockout plasmid (22). However, a recent genome-wide CRISPR/Cas9 knockout screen indicated that TLN4 does not contribute substantially to parasite fitness based on its phenotype score of 0.51 (on a scale of -7 to 3, with lower values indicating lower fitness) (23). TLN4 is a member of the M16 metalloproteinase subfamily of so-called “cryptases”, which are represented throughout the tree of life and include four TLN genes encoded in the *Toxoplasma* genome. M16 metalloproteinases are exemplified by the human Insulin-Degrading Enzyme (IDE) and feature a catalytic chamber (or crypt) that accommodates small polypeptides for degradation. Whereas characterization of TLN2 (TGME49_227948) and TLN3 (TGME49_257010) has not been reported, TLN1 (TGME49_269885) resides in the parasite secretory rhoptries where it plays an unknown role apart from contributing modestly to growth *in vitro* (24).

Here we show that TLN4 has the structural features of an active metalloproteinase and that it possesses proteolytic activity against a model polypeptide substrate. We also report that TLN4 deficient parasites show normal gliding motility, cell invasion, and replication, but have a delayed induced-egress phenotype. Our findings suggest that TLN4 contributes to *Toxoplasma* egress, identifying it as only the second microneme protein implicated in this event.

## RESULTS

### TLN4 has the structural features of an active metalloproteinase

TLN4 is a large (2,435 aa) protein that contains multiple domains including one active (A) and three inactive (IA) M16 proteinase domains, along with a long C-terminal extension that includes a repeat domain consisting of eight tandem repeats of a 28 aa sequence (Fig. 1A). Structural modeling of the M16 proteinase domains (residues 202-1367) using the human insulin degrading enzyme as a template suggests that each domain forms a similar αβ roll fold, with the domains arranged around a central chamber (Fig. 1B), as expected for an M16 family metalloproteinase (25). When viewed from a perspective of inside the chamber or “crypt”, a putative active site is visible (Fig. 1C) with the characteristic HXXEHX_N_EX_6_E binding motif for catalytic Zn^2+^ arranged on two α-helices (Fig. 1D) that are separated by 60 amino acids in TLN4 (Fig. 1E). These features are consistent with TLN4 being an active metalloproteinase of the M16 family.

**Figure 1.**
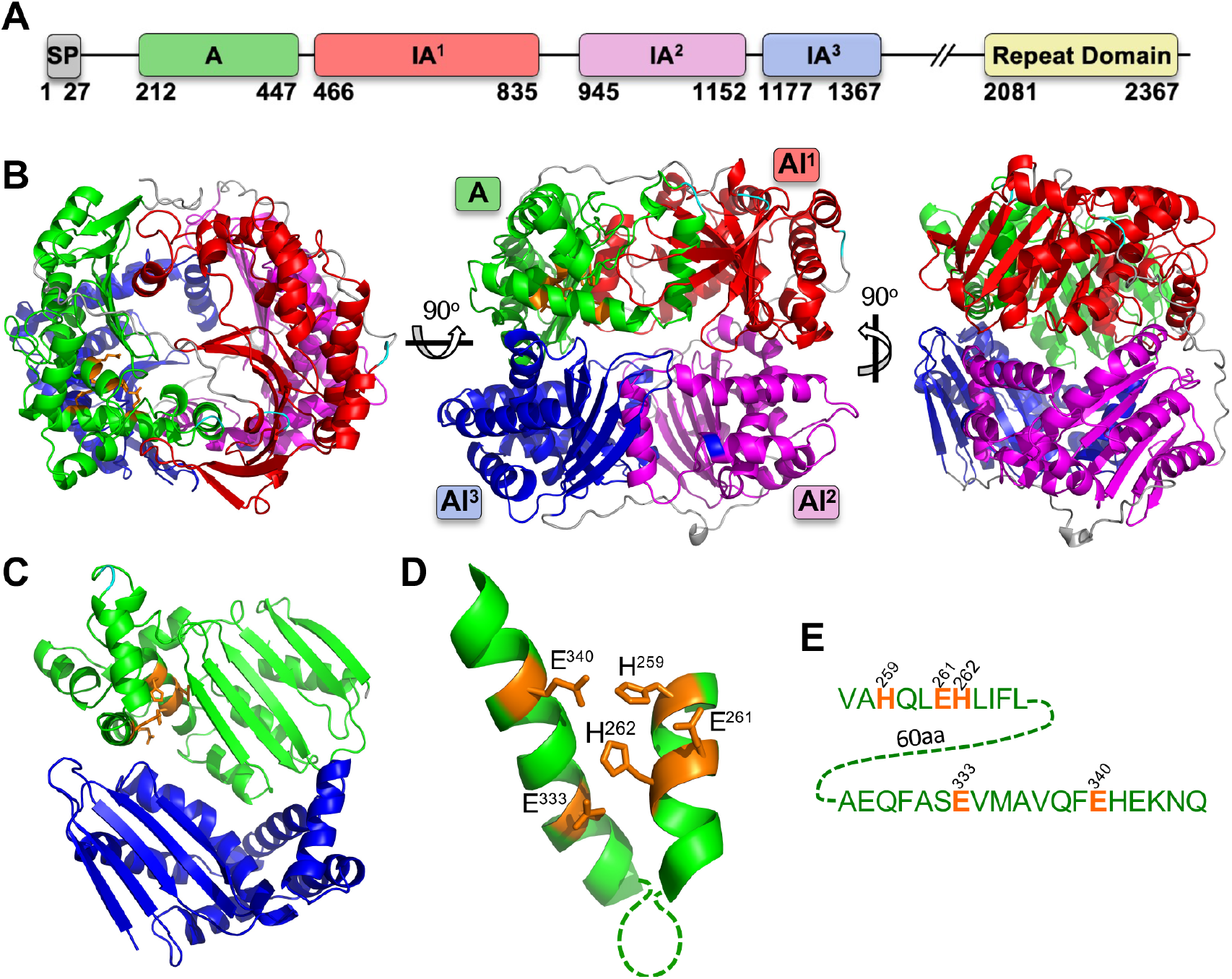
Structural features of TLN4. A) Domain structure of TLN4 illustrating its possession of a signal peptide (SP), and based on the IDE structure, a single active domain (A), 3 inactive domains (IA^1-3^), and a C-terminal repeat domain. Note that a largely featureless region between IA^3^ and the repeat domain is not shown and is instead denoted by //. Numbering indicates amino acid positions and is based on the sequenced cDNA of TLN4 (22). B) Structural model of TLN4 showing the arrangement of domains into a shell surrounding a central catalytic chamber. Three different views are provided. C) The A and AI^3^ domains viewed from the center of the chamber with resides that coordinate zinc binding shown in orange. D) Zoom in of the adjacent alpha helices of the A domain showing the residues (orange) that are predicted to coordinate binding to zinc. E) Sequence of the alpha helices shown in D) including residues (orange) that likely mediate binding to zinc as a cofactor for proteolytic activity.

### Recombinant TLN4 is capable of processing β-insulin

M16 family proteinases typically act upon peptides and polypeptides that are sufficiently small to fit in the crypt. To determine if TLN4 possesses proteolytic activity, we expressed and purified a recombinant form of TLN4 lacking the C-terminal extension. TLN4_209 – 1295_ was expressed using a bacterial expression system and was purified and refolded. The purified protein migrated on SDS PAGE as a prominent 130 kDa band and behaved as a monodispersed protein when analyzed by size exclusion chromatography (Fig. 2A). Upon incubating recombinant TLN4 with β-insulin as a model substrate, we observed the generation of several cleavage products (Fig. 2B), which were confirmed by mass spectrometry to originate from β-insulin (Fig. 2C). Mapping of the cleavage sites on the sequence of β-insulin revealed a preference for TLN4 cleavage in the central region of the polypeptide but no obvious preference for recognition of specific amino acids. Collectively our findings suggest that TLN4 is an active protease that is capable of processing a model substrate.

**Figure 2.**
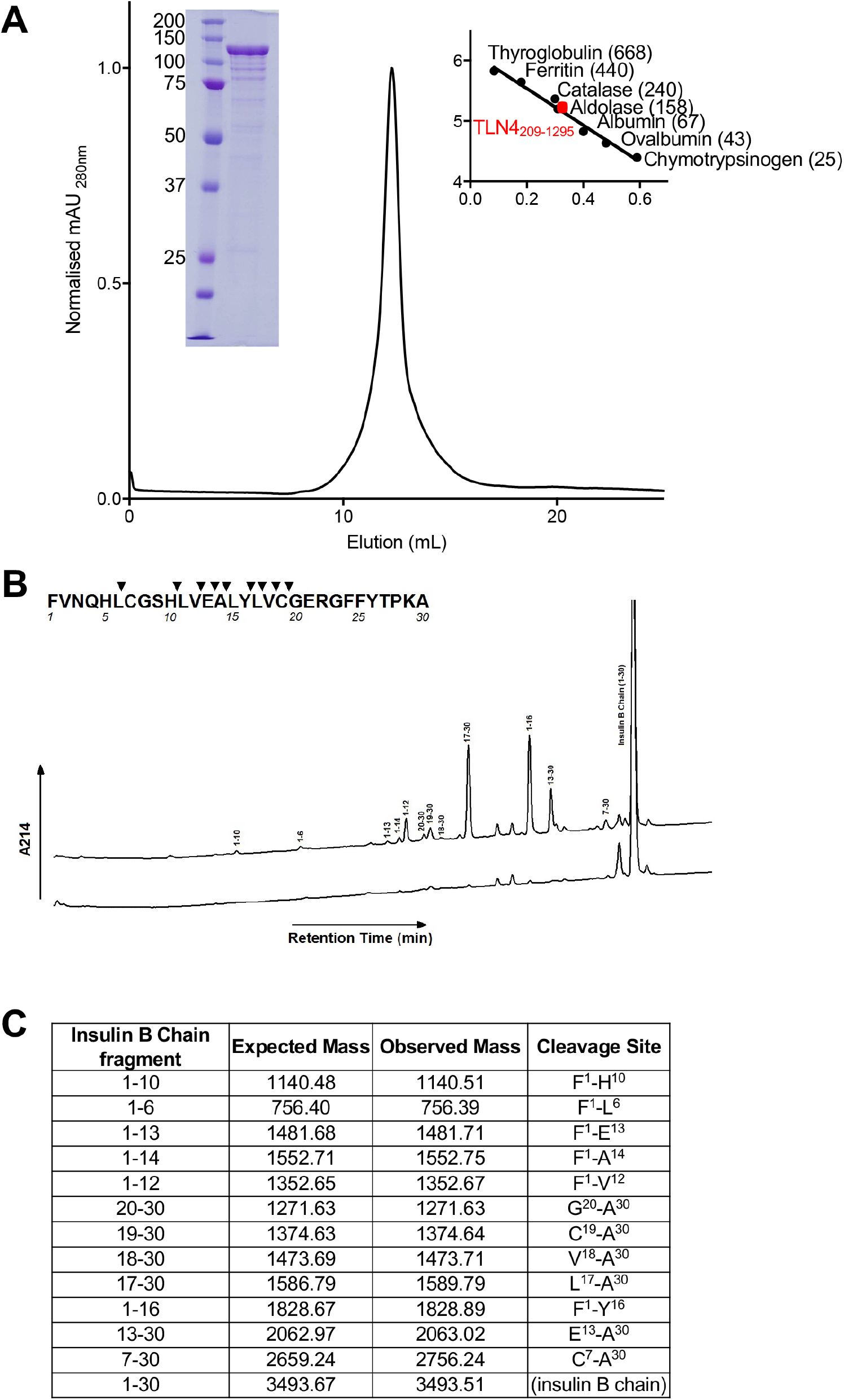
Recombinant TLN4 proteolytic activity using β-insulin as a model substrate. A) Chromatogram showing the elution of recombinant TLN4 as a monodispersed peak from size exclusion chromatography. SDS-PAGE of purified TLN4 is shown inset on the left, with molecular weight markers indicated on the right of elution peak. Values in brackets indicate molecular weight in kDa. B) HPLC profile of insulin B chain cleaved by TLN4. Hydrolysis products were determined by comparing a reaction allowed to incubate (top line) versus a reaction that was immediately stopped (bottom line). Captured peaks are labeled with their residue number and cleavage map was developed (inset). C) Mass spectrometry analysis of captured hydrolysis products from Insulin B Chain are shown with their expected mass and observed mass along with the fragment and cleavage sites that each product represents.

### *TLN4* is amenable to genetic disruption

In a previous study of TLN4, it was proposed that TLN4 contributes to parasite fitness because TLN4-knockout parasites could be detected in a population of transfected RH parasites but were lost through further culturing (22). To enhance the recovery of knockout parasites we utilized RHΔ*ku80*Δ*hxg* parasites (WT hereafter) and transfected them with a hypoxanthine xanthine guanine phosphoribosyl transferase (HXGPRT) selectable cassette flanked by 5’ and 3’ *TLN4* homology regions (Fig. 3A). Individual knock-out parasite clones were tested by PCR for integration of the selectable marker at the 5’ end (Fig. 3B, left gel), at the 3’ end (Fig 3B, middle gel), and for the absence of the TLN4 gene (Δ*tln4*) (Fig 3B, right gel). We then genetically complemented Δ*tln4* parasites by expression of the *TNL4* cDNA containing two copies of an HA epitope tag inserted at aa position 847 after the first inactive domain (TLN4-IA^1^-HA_2_) to generate Δ*tln4TLN4* parasites, which were confirmed by PCR (Fig. 3C). The difference in fragment size in the WT and the Δ*tln4TLN4* is due to the presence of introns in the WT but not in the cDNA of the complement strain. Expression of TLN4 in Δ*tln4TLN4* parasites was confirmed by fluorescence microscopy (Fig. 3D) and by western blot, with the expected shift in migration of the ~33 kDa processed species due to insertion of HA (Fig. 3E). The loss of the TLN4 protein in Δ*tln4* was also confirmed by western blot (Fig. 3E).

**Figure 3.**
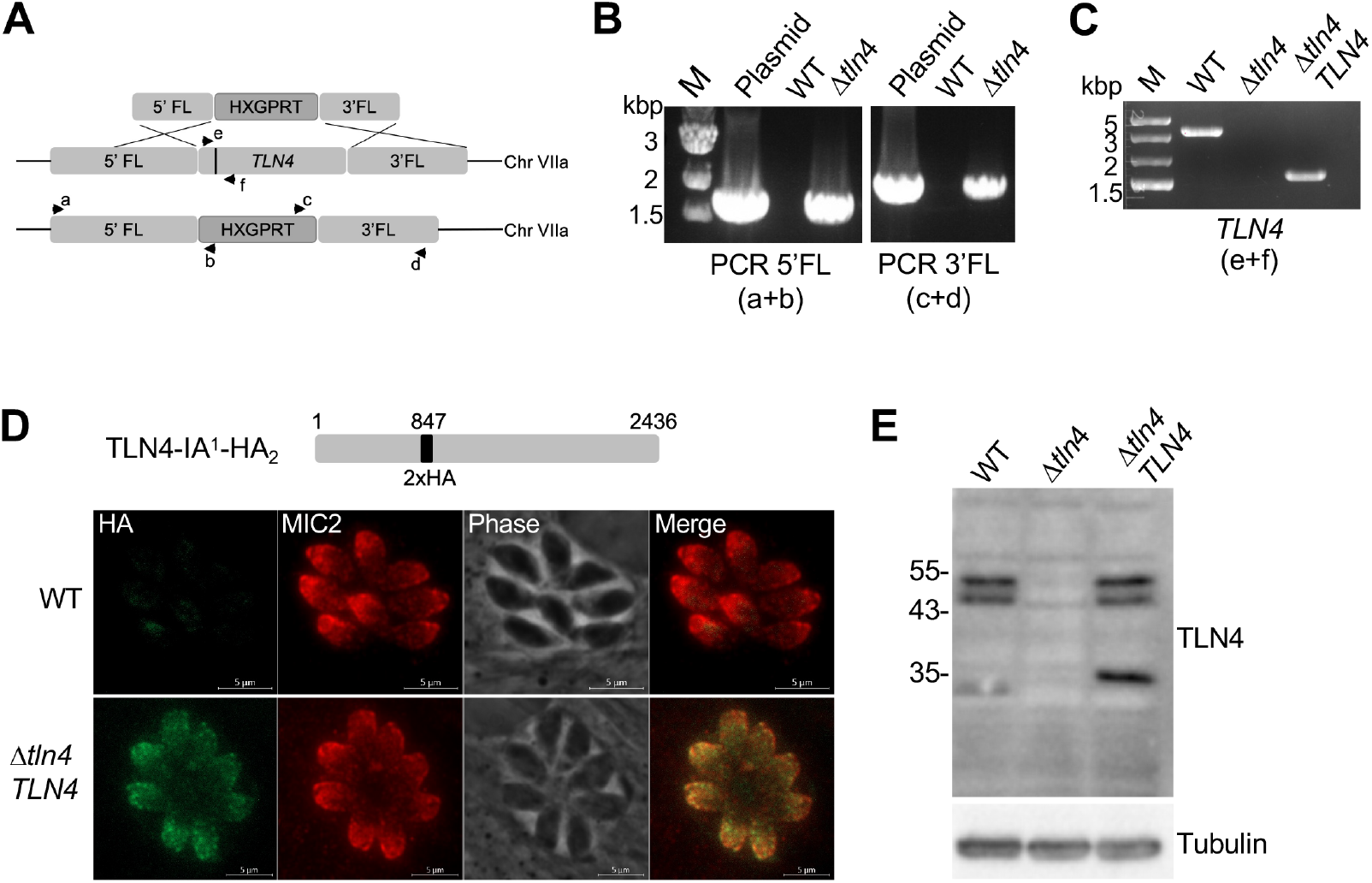
Genetic disruption of *TLN4*. A) Schematic showing the TLN4 genomic locus and homologous recombination with a cassette containing a *TLN4* 5’flank, the HXGPRT selectable marker, and a *TLN4* 3’flank. Arrows and letters indicate the placement of primers used in panel B, the black bar in TLN4 indicates an intron that permits distinction of the gene from the cDNA in panel C. Other introns of TLN4 are not shown. B) PCR validation of selectable marker integration at the 5’ (primers a+b) and 3’ (primers c+d) end. M, molecular weight marker. C) PCR validation of *TLN4* deletion (Δ*tln4*) and complementation (Δ*tln4TLN4*) with the cDNA (primers e+f). The differences in size are due to the presence of an intron in the WT genomic DNA versus the cDNA in the complement strain lacking the intron. D) Schematic showing the complementation construct with a 2xHA tag in the full-length TLN4 coding sequence. Immunofluorescence panel shows the micronemal localization of the complemented strain, co-stained with anti-MIC2. E) Mouse antibodies to TLN4 detect the ~55 kDa doublet and the ~32 kDa of TLN4 in the lysates of WT and the Δ*tln4TLN4*. Slower mobility of the smallest band in the complement strain is due to the HA tag. Sizes (in kiloDaltons) of molecular weight markers are shown to the left. Detection of *T. gondii* tubulin was included as a loading control.

### TLN4 does not contribute to invasion, replication, gliding motility, or virulence

To assess whether TLN4 plays a role in the lytic cycle, plaque assays were performed. Compared to parental parasites, Δ*tln*4 parasites showed significantly smaller plaques (Fig. 4A,B), suggesting a defect in one or more steps in the parasite lytic cycle. This phenotype that was fully restored in the complement strain. We next tested each step in the lytic cycle to identify the basis of smaller plaques. Parasites deficient in TLN4 showed normal invasion (Fig. 4C), replication at 17 h and 26 h post-infection (Fig. 4D) and all types of gliding motility were observed (Fig. 4E). Finally, no significant differences were observed in mouse survival of acute infection upon intraperitoneal infection with 10 or 100 tachyzoites (Fig. 4F).

**Figure 4.**
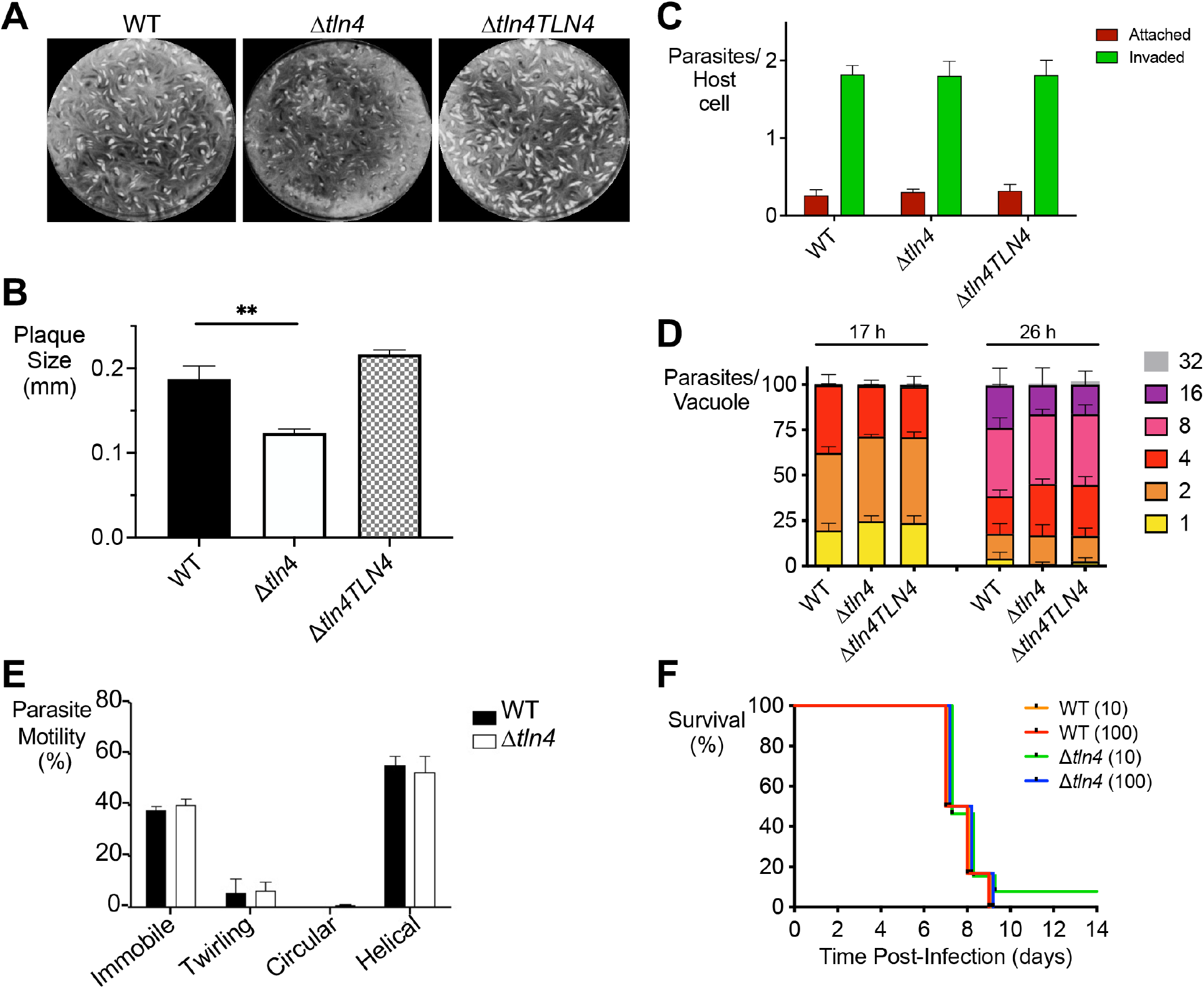
Effect of TLN4 deficient parasites on the lytic cycle. A) Plaque assays show smaller plaques in Δ*tln4* parasites. One hundred parasites of each strain were inoculated into 6-well plates and allowed to grow for 7 days undisturbed. Wells were stained with crystal violet. B) Individual plaque sizes were measured using ImageJ. Data represent three independent experiments with triplicates within each experiment. C) Parasites lacking TLN4 show normal invasion. HFF cells in 8-well chamber slides were inoculated with parental, Δ*tln4*, or Δ*tln4TLN4* parasites and allowed to invade for 20 min prior to fixation. Wells were differentially stained with SAG1 antibodies to detect attached or invaded parasites per host cell nucleus. D) Parasites lacking TLN4 replicate normally. Parasites were inoculated into 8-well chamber slides and allowed to replicate for 17 h or 26 h prior to fixation and enumeration of parasites per vacuole. Data in invasion and replication graphs represent means ± SEM of three independent experiments each with triplicate samples. E) Δt*ln4* parasites show normal modes of gliding motility. Video microscopy to enumerate the types of gliding motility show the percentages of immobile parasites, and parasites performing twirling, circular, and helical gliding in all strains tested. Graphs indicate the means ± SEM of three independent experiments. * *p* 0.05 by Student’s *t*-test. F) TLN4 does not play a role in acute virulence. Swiss-Webster mice were infected intraperitoneally with 10 or 100 tachyzoites of WT or Δ*tln4*, and survival time was enumerated; 12 mice were infected for each strain and inocula.

### Δ*tln4* parasites are defective in efficient egress

The last step of the lytic cycle is egress of the intracellular parasites from host cells, which can be induced by addition of the calcium ionophore A23187. Whereas most wild-type parasites egressed from host cells within 2 min of ionophore addition, significantly fewer Δ*tln4* parasites egressed in the allotted time (Fig. 5A). This phenotype is mostly reversed in Δ*tln4TLN4* parasites. The deficiency in egress was more pronounced when zaprinast, a phosphodiesterase inhibitor, was used as an inducer (Fig. 5B), which may be due to its mode of action. Whereas A23187 overtly elevates cytosolic calcium by mobilizing it from intracellular stores and the medium, zaprinast treatment triggers the release of calcium from intracellular stores via activation of protein kinase G and inositol triphosphate signaling (4, 26). A time-course of egress further illustrated the delay in Δ*tln4* parasites, with a statistically significant difference in the time to 50% egress compared to WT parasites (Fig. 5C). Δ*plp1* parasites, which were included as a reference strain, showed an expected severe impaired in egress. An alternative and complementary measure of parasite egress is the release of host lactate dehydrogenase (LDH) into the surrounding media due to loss of host cell integrity. After normalizing LDH release to WT parasites, Δ*tln*4 parasites showed ~45% release of LDH, while the complement strain nearly restored LDH release to wild-type levels (Fig. 5D). In examining the excreted-secreted-antigen fraction of extracellular parasites, conducted in buffers with neutral or acidic pH (to mimic PV acidification; (13–15)), there was no change in the processing of PLP1 in the Δ*tln4* parasites compared to WT parasites (Fig. 5E). Together these findings suggest a role for TLN4 in *T. gondii* egress from host cells that is independent of PLP1 processing.

**Figure 5.**
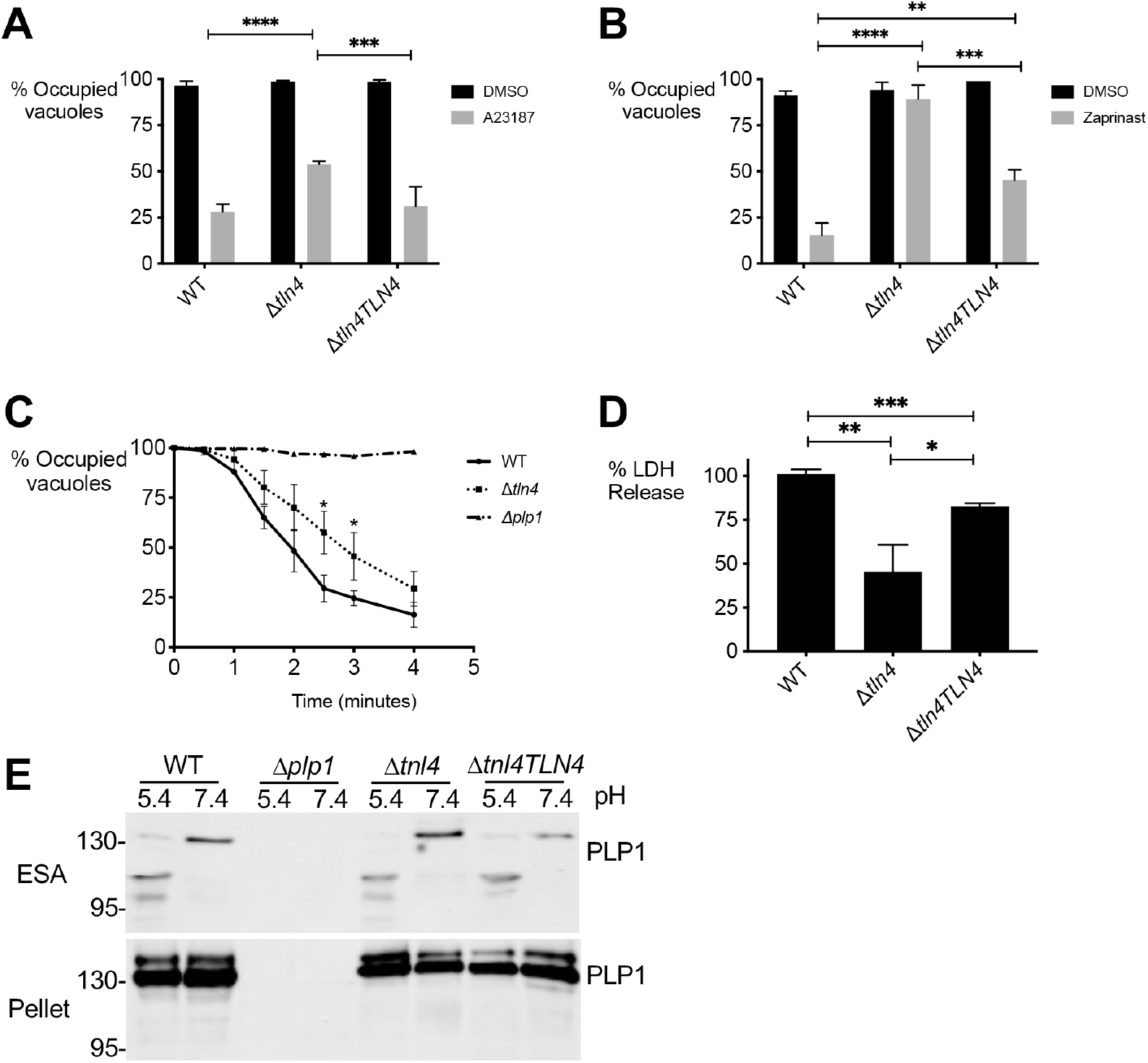
TLN4 deficient parasites are defective in egress. A) and B) Parasite egress was quantified by immunofluorescence microscopy. Thirty-hour vacuoles were treated with DMSO or 2 μM A23187 (A) or 200 μM zaprinast (B) for 5 min prior to fixation and enumeration. \**p*≤0.05, **p≤0.01, ***p≤0.001, \*\*\*\**p*≤0.0001 by Student’s *t*-test. C) Time-course of egress with 2 μM A23187, quantified by immunofluorescence as in A and B. Δ*plp1* parasites were included as a negative control for egress. D) Lactate dehydrogenase (LDH) release following induction with 100 μM zaprinast was used as a measure of egress. Data are normalized to WT LDH release. All graphs shown are means ± SEM from three biological replicates each with triplicate samples. E) Parasites lacking TLN4 has normal PLP1 processing. Pellet and ESA were collected from parasites at pH 5.4 and 7.4, separated on SDS-PAGE, blotted onto membranes, and probed with rabbit anti-PLP1 antibodies. Processing of PLP1 in the ESA of all strains expressing PLP1 observed at pH 5.4. The pellet blot (bottom) acts as a loading control.

## DISCUSSION

TLN4 was previously shown to be an extensively processed microneme protein (22), but its contribution to the lytic cycle was not determined due to the inability to generate a knockout of this gene in the wildtype RH strain background. In this study we were able to isolate a TLN4 knockout strain by utilizing the more genetically tractable RHΔ*ku80* strain. Plaque assays showed that Δ*tln4* parasites affected the lytic cycle, based on the smaller plaque sizes observed. Assessing the known steps of the lytic cycle showed that there were no defects in invasion, replication, or types of gliding motility performed by the parasites, nor was there any effect on parasite virulence. The only deficiency observed was in induced egress, assessed by a static egress assay, a time-course of egress, and LDH-release. Complementation of Δ*tln4* with a TLN4 construct under the control of the endogenous TLN4 promoter fully restored the plaque defect and largely restored the egress deficiency.

*In vitro* investigation into TLN4 suggests that the protein is an active protease based on cleavage of β-insulin as a model substrate, supporting a potential proteolytic function within the parasite. Homology modeling indicates that TLN4 likely adopts the typical M16 structure and contains a catalytic chamber or “crypt”, which is one of the defining features of the M16 family of metalloproteinases. Studies with other M16 enzymes indicate that the crypt encapsulates and cleaves polypeptides, with the substrate being determined by the size and charge of the crypt as well as the flexibility of the substrate(s) (27, 28). Although it is possible that TLN4 plays a direct role in egress by e.g., facilitating the disruption of the PVM, an indirect contribution of TLN4 to egress is also plausible. An indirect role could involve proteolytically activating another parasite protein that contributes to egress or degrading a protein that would otherwise compromise egress. Identifying substrates of TLN4 in future studies might reveal the basis for its contribution to parasite exit from host cells.

PLP1 is the only microneme protein identified to date to have a direct role in egress. Interestingly, mice infected with Δ*tln4* have no virulence defect, whereas Δ*plp1* parasites are markedly virulence attenuated. This notable distinction could be because the egress defect of Δ*tln4* parasites is less pronounced than that for Δ*plp1* or it could be due to additional roles for PLP1 during infection of mice. Future studies identifying other secretory proteins that contribute to egress to varying degrees along with other work defining how PLP1 shapes the outcome of infection should help distinguish between these possibilities.

Among the Apicomplexa, the M16A subfamily is constrained to coccidian parasites, including close (e.g., *Hammondia hammondia*, *Neospora caninum*) and more distant (*Sarcocystis neurona*, *Eimeria* spp., *Cryptosporidium* spp.) relatives of *T. gondii* (orthomcl.org). Whereas the genome of *T. gondii* encodes 11 M16 metalloproteinases, of which 4 belong to the M16A subfamily (29), the genome of the intestinal parasite *Cryptosporidium parvum* encodes an expanded family of 22 M16 metalloproteinases, 18 of which are M16A members. Among these, INS1 was recently shown to be necessary for formation of macrogamonts, the female sexual stage of *C. parvum* (30). The results of reciprocal BLAST searches indicate that TLN4 is most closely related to *C. parvum* INS-15 (cgd3_4260) and INS-16 (cgd3_4270), which are closely related orthologs (83% identical) (30, 31). INS-15 localizes to the mid-apical region of *C. parvum* sporozoites and possibly merozoites (31). Antibodies to INS-15 impaired sporozoite cell invasion, suggesting a role in parasite entry into host cells. Future studies involving the targeted deletion of M16A family members in *C. parvum* and other coccidian parasites will be necessary to appreciate further their contributions to parasite infection biology.

## METHODS AND MATERIALS

### Structural modeling of TLN4

The full amino acid sequence of ME49 strain TLN4 was used to query Phyre2, which identified human insulin degrading enzyme (PDB 2JBU) as the top scoring template. This analysis resulted in a structural model of TLN4 encompassing amino acids 202-1367. The structural model was visualized using PyMOL v2.3.2.

### Expression and purification of recombinant TLN4

A region encoding domains A to IA-3 of the *TLN4* cDNA from RH strain (amino acids 209 – 1295) was amplified with TLN4.625.NdeI.F (5’-GGAATTCCATATGAGAGACACGAGCGCGTACTCGG-3’) and TLN4.3885.Not.R (5’-AAGGAAAAAAGCGGCCGCGCTAAGCCACCGGAGGCGCTCGAGGAAAGC-3’) primers, and cloned into the bacterial expression vector pET21a, introducing an in-frame C-terminal hexa-Histidine tag to the construct. The *E. coli* expression construct containing the *TLN_4209-1295_* ORF was verified by DNA sequencing.

The *TLN4_209-1295_* expression plasmid was transformed into *Escherichia coli* BL21 Gold cells selected on 2x yeast-tryptone (2YT) broth-agar plates containing 100 μg/mL ampicillin. 20 mL of 2YT supplemented with ampicillin was grown to confluence overnight and then added to 1 L of 2YT containing ampicillin and incubated with shaking at 37°C until an OD_600_ of ~ 0.85 was reached. Cultures were then induced with a final concentration of 1 mM IPTG overnight at 16°C.

Harvest and cell lysis of the overnight expression culture showed that TLN4_209-1295_ expressed to high yields but was insoluble. Inclusion bodies were prepared by harvest of the insoluble fraction of the whole cell lysate and resuspension with 30 mL of wash buffer I (25 mM Tris pH 8.0, 300 mM NaCl, 1% Triton X-100). The suspension was homogenized until an even consistency was obtained. The inclusions bodies were then washed twice by centrifugation and homogenization in wash buffer I. Three subsequent washes with wash buffer II (25 mM Tris pH 8.0, 300 mM NaCl) were performed to remove the excess Triton X-100. Purified inclusion bodies were stored at −80°C until required. Proteins were resolubilized in 25 mM Tris pH 8.0, 250 mM NaCl, 6 M guanidine hydrochloride, 2 mM beta-mercaptoethanol by rotation at 4°C before centrifugation. Harvest supernatant was applied to Ni-NTA resin and purified via metal affinity chromatography. Purity of the denatured purified protein was assessed via SDS-PAGE and western blot.

Denatured TLN4_209-1295_ was diluted to a final concentration < 1 mg/mL and applied dropwise to 100 mL of refold buffer (50 mM Tris pH 8.2, 250 mM NaCl, 0.5 M arginine, 0.44 M sucrose, 4 mM reduced glutathione, 0.4 mM oxidised glutathione, 0.05 mM Tween-20) that was constantly stirring at 4°C. Refolding proceeded for 4 h at 4°C with gentle stirring. The refolded protein was then dialyzed overnight against 25 mM Tris pH 8.0, 300 mM NaCl at 4°C. Post-dialysis, the protein was concentrated to approximately 0.5 mL before application to a Superdex 200 10/300 column for further purification via size-exclusion chromatography.

### TLN4 proteolytic activity

Hydrolysis of Insulin B Chain was measured via reverse phase high performance liquid chromatography (HPLC) following incubation of TLN4 (160 μg/mL) with insulin B chain (42 μM, Sigma Aldrich) in 10 mM Tris for 88 h. HPLC was carried out in a Vydac C4 HPLC column using a linear gradient from 0.1% trifluorocacetic acid (TFA) in 95% water, 5% acetronitrile to 0.1% TFA in 50% acetonitrile/water at a flow rate of 1 mL/min. Hydrolysis products were detected at 214 nm and collected for analysis. The obtained peptide products were analyzed on an Applied Biosystems 4800 MALDI TOF/TOF Proteomics Analyzer at the University of Kentucky Proteomics core.

### Parasite culture, transfection, and selection

*T. gondii* tachyzoites were maintained by growth on monolayers of human foreskin fibroblasts (HFF) in Dulbecco’s Modified Eagles Medium (DMEM) containing 10% Cosmic Calf Serum (GIBCO), 2 mM Glutamine, 10 mM HEPES, 50 μg/mL Penicillin/Streptomycin (D10 Complete). To generate the TLN4 KO, a knockout construct (described in (Laliberte & Carruthers, 2011)) containing 1.5 kb of TLN4 5’ and 3’ flanking regions was used and the HXGPRT selectable marker was replaced with the DHFR-TS selectable marker cassette. This construct was transfected into RHΔ*ku80*Δ*hxg* parasites with a BioRad Gene Pulsar II with 1.5 kV voltage, 25 μF capacitance and no resistance setting.

Pyrimethamine selection was applied the day after transfection and clones were isolated by limited dilution in 96-well plates. Knockout clones were tested for proper 5’ integration (a) 5’-agttgcagccagaggcagaagcaagtcc-3’ and (b)

5’-cagtcagataacaggtgtagcg-3’ and for proper 3’ integration (c) 5’-gcgggtgacgcagatgtgcgtgtatcc-3’; (d) 5_-gaaaagtgtctgcgtgttagcagc-3. Knockout clones that were complemented with the TLN4.HA construct were identified with primers TLN4.322.F (e) 5’GGCTTTTCTGCTTCGTCAAC-3’ and TLN4.1910.R (f) (5’AAGAGCAGTGGGCTGAAAAA-3’).

### Invasion assays

Invasion assays were performed as previously described (32). Briefly, 1×10^7^ parasites were used to infect sub-confluent HFF monolayers in 8-well chamber slides for 20 min before fixation with paraformaldehyde. Slides were differentially stained with anti-SAG1 antibodies to differentiate attached versus invaded parasites.

### Egress assays

Induced egress was performed as described previously (15). Briefly, parasites grown in HFFs in an 8-well chamber slide for 30 h were treated with 1% dimethyl sulfoxide, 2 μM A23187, or 200 μM zaprinast in assay buffer (Hanks’ buffered salt solution containing 1 mM CaCl_2_, 1 mM MgCl_2_, and 10 mM HEPES) and incubated for 2 min in a 37°C water bath. Egress was stopped by addition of 2x fixative (8% formaldehyde in 1x PBS). Immunofluorescence was performed with rabbit anti-SAG1 to identify parasites and mouse anti-GRA7 (a generous gift from Peter Bradley) to identify the parasitophorous vacuole membrane. At least 10 fields of view (400x total magnification) per condition were enumerated as occupied or unoccupied.

LDH egress assays were performed as previously described (20). Briefly, parasites were grown in HFF monolayers in 96-well plates for 30 h. Wells were washed with Ringer’s buffer and then treated with 100 μM zaprinast diluted in Ringer’s buffer. Plates were incubated at 37°C for 20 min before removal and placement on ice. Fifty microliters of the supernatant were removed and centrifuged in a separate round-bottom plate. Thirty microliters of supernatant was subsequently removed and release of lactate dehydrogenase was determined using an LDH Cytotoxicity Colorimetric Assay kit (BioVision).

Time-course egress assays were performed as described above, with time points of 0.5, 1, 1.5, 2, 2.5, 3 and 4 min following induction with A23187, at which time wells were fixed, and IFA performed with rabbit anti-SAG1 and mouse anti-GRA7.

### Plaque and replication

Parasites were inoculated into wells of a 6-well plate and allowed to replicate undisturbed for 7 days. The wells were then stained with 0.2% Crystal violet for 5 min and rinsed with ddH_2_O. Images of the wells were scanned, and plaque number and size were analyzed with Image J.

For the replication assay, cells of an 8-well chamber slide were inoculated with 1.25×10^5^ tachyzoites and allowed to invade and grow for 18 h prior to fixation and indirect immunofluorescence.

### Live gliding video microscopy

Glass-well dishes (MatTek) were coated with 50%FBS/50%PBS overnight at 4°C or 1 h at 37°C and washed with PBS before use. Parasites were filter-purified and resuspended in HHE (Hank’s salt solution, 10mM HEPES, EDTA). Parasites were then added to coated dishes and allowed to settle for 5 min at room temperature. Dishes were then moved to a 37°C chamber with 5% CO_2_ and warmed for 5 min prior to starting video recording. Each video consisted of 1 frame/sec recorded for 90 sec. Enumeration of the types of gliding motility were carried out by examining the videos in addition to maximum projection images generated by the Zeiss AxioVision software.

### Mouse infections

All laboratory animal work in this study was carried out in accordance with policies and guidelines specified by the Office of Laboratory Animal Welfare, the US Department of Agriculture, and the American Association for Accreditation of Laboratory Animal Care (AAALAC). The University of Michigan Committee on the Use and Care of Animals (IACUC) approved the animal protocol used for this study (Animal Welfare Assurance A3114–01, protocol PRO00008638). Freshly egressed parasites were filter purified in PBS, washed, counted, and injected intraperitoneally in 200 μl of PBS into 6-8-week-old female Swiss Webster mice. The experiment was performed once with 4 groups of 12 mice each infected with 10 or 100 tachyzoites of WT or Δ*tln4*. Mice were given water and food ad libitum, monitored twice daily and were humanely euthanized upon showing signs of moribundity.

### Western blotting

Parasite lysates were generated by filter-purifying parasites followed by centrifugation, 1x wash with cold PBS, and resuspension in >90°C 1x sample buffer. Lysates were boiled for 5 min prior to running on SDS-PAGE gels. Gels were semi-dry electroblotted (Bio-Rad) onto polyvinylidene fluoride membranes and sequentially probed with mouse anti-TLN4 (22) and horse radish peroxidase conjugated goat anti-mouse (Jackson ImmunoResearch). Bands were revealed by enhanced chemiluminescence with SuperSignal West Pico PLUS Chemiluminescent Substrate (Thermo Fisher Scientific) and documented with a Syngene Pxi imaging system.

### PLP1 processing

Extracellular parasites (Δ*ku80*Δ*hxg (*WT), Δ*plp1*, Δ*tln4*, and Δ*tln4TLN4*) were resuspended in PBS in either pH 5.4 or 7.4 and ESA was collected. Pellet lysates and ESA were separated on SDS-PAGE gels and membranes were probed with antibodies against rabbit anti-PLP1.

## ACKNOWLEDGEMENTS

This work was supported by National Institutes of Health grant R01AI46675 (VBC) and 1R01GM130954 and 5P30GM110787 (LBH). AJS received fellowship support from the American Heart Association. We thank Tracey L. Schultz for technical assistance.

